# Random noise promotes slow heterogeneous synaptic dynamics important for robust working memory computation

**DOI:** 10.1101/2022.10.14.512301

**Authors:** Nuttida Rungratsameetaweemana, Robert Kim, Thiparat Chotibut, Terrence J. Sejnowski

**Affiliations:** Department of Biomedical Engineering, Columbia University, New York, NY 10027, USA; Computational Neurobiology Laboratory, Salk Institute for Biological Studies, La Jolla, CA 92037, USA; Neurology Department, Cedars-Sinai Medical Center, Los Angeles, CA 90048, USA; Chula Intelligent and Complex Systems, Department of Physics, Chulalongkorn University, Bangkok, Thailand; Institute for Neural Computation, University of California San Diego, La Jolla, CA 92093, USA; Division of Biological Sciences, University of California San Diego, La Jolla, CA 92093, USA

**Author notes:** Correspondence (T.C.) and (T.J.S.). Equal contribution (Nuttida Rungratsameetaweemana, Robert Kim).

## Abstract

Recurrent neural networks (RNNs) based on model neurons that communicate via continuous signals have been widely used to study how cortical neurons perform cognitive tasks. Training such networks to perform tasks that require information maintenance over a brief period (i.e., working memory tasks) remains a challenge. Critically, the training process becomes difficult when the synaptic decay time constant is not fixed to a large constant number for all the model neurons. Here, we show that introducing random noise to the RNNs not only speeds up the training but also produces stable models that can maintain information longer than the RNNs trained without internal noise. Importantly, this robust working memory performance induced by internal noise during training is attributed to an increase in synaptic decay time constants of a distinct subset of inhibitory units, resulting in slower decay of stimulus-specific activity critical for memory maintenance.

## Introduction

It is widely acknowledged that the cortex exhibits a high level of spontaneous activity that appears unrelated to task-specific neural codes or behaviors. However, recent works have demonstrated that such “cortical noise” contains information about the environmental context and has a direct impact on downstream behavioral outcomes [1–3]. For instance, Musall et al. [1] showed that the cortical noise in mice contains information about the visual stimulus even in the absence of a task, suggesting that it may play a role in sensory processing. Similarly, Stringer et al. [3] found that the cortical noise in mice contains information about the animal’s location and movement speed, which is crucial for navigation. Furthermore, previous studies have also shed light on the significance and relevance of cortical noise to cognitive processes. For example, Caron et al. [4] showed that the random structures of the olfactory system in *Drosophila* optimized the diversity of odor representations in neural circuits. Together, these findings challenge the traditional view of cortical noise as mere “background noise,” highlighting its potential role in cognitive functions.

In addition to the experimental findings, there is growing evidence from computational and modeling studies that introducing noise during the training process can lead to improved stability and robustness of neural networks. Specifically, several studies have demonstrated that injecting Gaussian noise during the training process of multi-layer perceptron (MLP) and recurrent neural networks (RNNs) can improve their performance [5–7]. For example, Lim et al. [7] examined the impact of injecting noise into the hidden states of vanilla RNNs and found that it contributed to stochastic stabilization through implicit regularization [8]. Additionally, Camuto et al. [6] studied the regularization effects induced by Gaussian noise in MLPs and showed that the explicit regularization provided several benefits, including increased robustness to perturbations.

Despite the demonstrated benefits of noise injection in vanilla RNNs and MLPs, it is not yet clear whether these findings extend to more biologically plausible RNNs that incorporate neuronal firing rate dynamics. It is also unclear if introducing noise can improve the cognitive capabilities of these RNNs. We hypothesize that incorporating noise into such biologically plausible RNNs will give rise to persistent activity, which in turn will be crucial for enhancing working memory performance.

In this study, we propose a systematic approach to address these questions. Specifically, we investigate the impact of noise during training of firing-rate RNNs to perform tasks that require different cognitive functions, such as decision making and working memory. We show that the introduction of noise during training significantly enhances the RNN performance on tasks that specifically require working memory. By dissecting the networks trained with noise and employing stability analysis methods, we further show that noise induces slow dynamics in inhibitory units and forces these units to be more active, resulting in more stable memory maintenance. These findings aligned with recent experimental and theoretical studies that place specific subtypes of inhibitory neurons at the center of working memory computations [9–13]. Therefore, our study illustrates how seemingly random noise in the cortex could lead to specific changes in synaptic dynamics critical for complex cognitive functioning.

## Results

### Biologically plausible RNN model and task overview

Even though recent advances in deep learning and artificial intelligence (AI) have greatly increased the functionality and capability of artificial neural network models, it is still challenging to train a network of model neurons to perform cognitive tasks that require memory maintenance. Models based on recurrent neural networks (RNNs) of continuous-variable firing rate units have been widely used to reproduce previously observed experimental findings and to explore neural dynamics associated with cognitive functions including working memory, an ability to maintain information over a brief period [14–17].

We study the following RNN model composed of excitatory and inhibitory rate units:

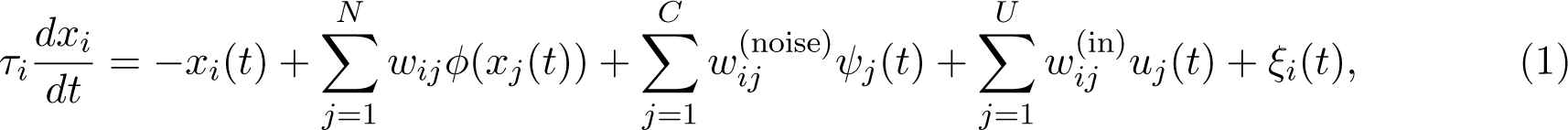

where τ*_i_* and *x_i_* refer to the synaptic decay time-constant and synaptic current variable, respectively, for unit i. The synaptic current variable is converted to the firing-rate estimate via a nonlinear transfer function (ϕ(*·*)). Throughout this study, we employed the standard sigmoid function for ϕ. w*_ij_* is the synaptic strength from unit *j* to unit *i*, and *u*(*t*) is the task-specific input data given to the network via *w*^(in)^ (see *Methods*). Each neuron in the model received “sensory” noise (ξ(*t*)). In contrast to conventional rate-based RNN models, our model incorporated what we referred to as “internal” noise, where a set of independent noise signals sampled from a standard Gaussian distribution uncorrelated in time (ψ(*t*)) were introduced to the network through *w*^(noise)^ (see *Methods*).

The above firing-rate RNN model was trained using backpropagation through time (BPTT; [18]) to perform a task that involves maintaining information over a brief period (i.e., working memory task). The task is a delayed match-to-sample (DMS) task that requires the model to match the signs of the two sequential input stimuli (Fig. 1a; see *Methods*). While the model has shown success in various cognitive tasks [14–17], training the model with important biological constraints to perform the DMS task with a long delay period between the two input stimuli remains challenging. Notably, the training time increases exponentially as a function of the delay duration. As shown in Fig. 1b, the model required more trials to achieve successful training on the DMS task as the delay interval increased from 50 ms to 250 ms (all *P s* < 0.001, two-sided Wilcoxon rank-sum test). Moreover, when the synaptic decay time constants (τ) for all the units in the model were fixed at a small constant (20 ms), the training process failed to converge.

**Fig. 1:**
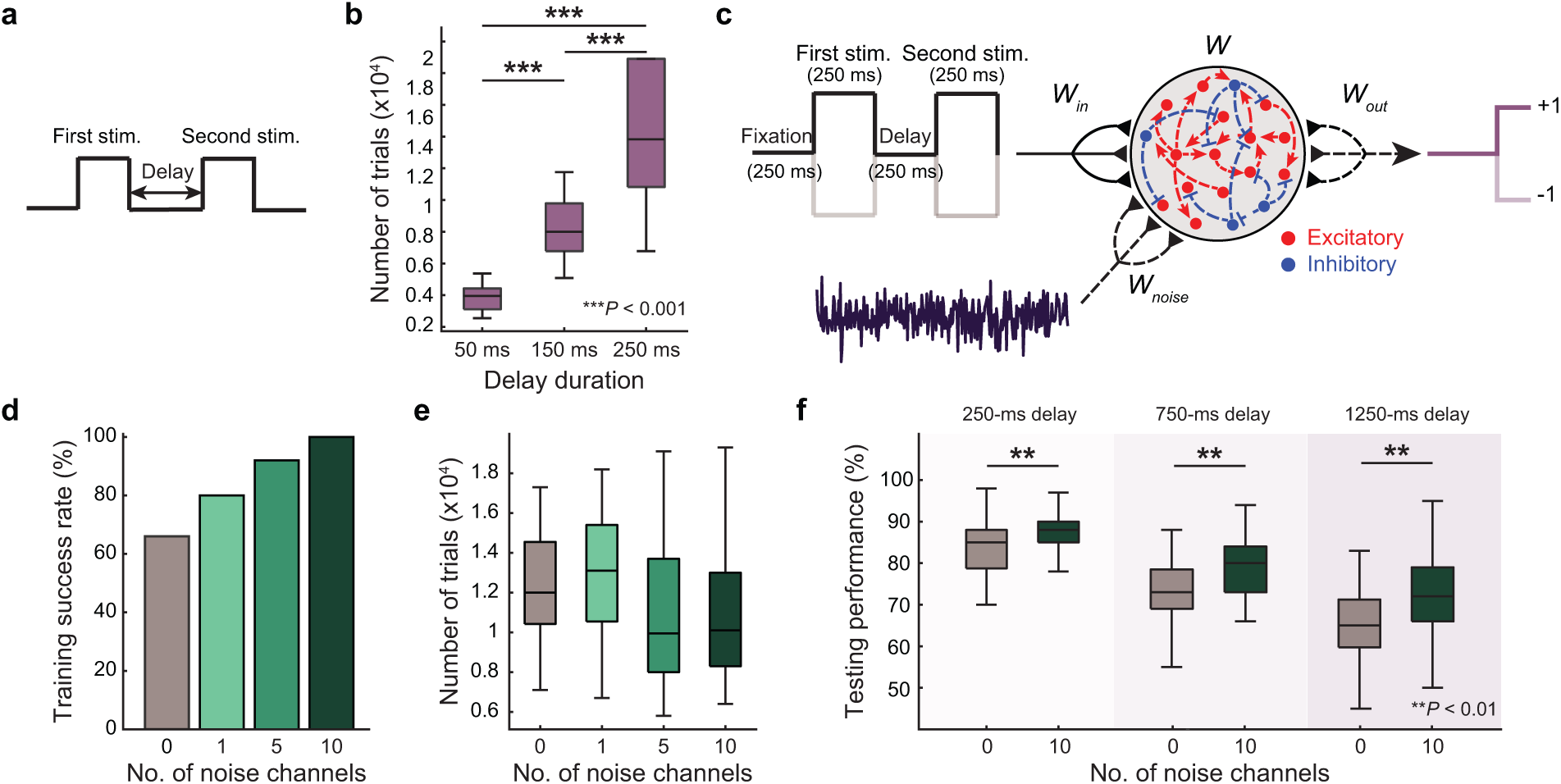
delayed match-to-sample (DMS) task and model schematic. **a**, A schematic diagram of a Delayed match-to-sample (DMS) task with two sequential stimuli separated by a delay interval. **b**, The number of trials/epochs needed to train continuous-variable RNNs increases exponentially as the delay interval increases. For each delay duration condition, we trained 50 firing-rate RNNs to perform the DMS task shown in **a**. The maximum number of trials/epochs was set to 20,000 trials for computational efficiency (all *P s* < 0.001, two-sided Wilcoxon rank-sum test). **c**, A schematic diagram illustrates the paradigm used to trained our RNN model on the DMS task in which one delay was present. We introduced and systematically varied the amount of noise in the RNN network to study the effects of noise on memory maintenance in a biologically constrained neural network model. The model contained excitatory (red circles) and inhibitory (blue circles). The dashed lines represent connections that were optimized using backpropagation. **d**, Training performance of the RNN models on the DMS task. RNN models with varying amount of noise (i.e., 0, 1, 5, and 10 noise channels) were trained to perform this task. Training success rate was measured as the number of successfully trained RNNs (out of 50 RNNs). **e**, The average number of trials required to reach the training criteria. **f**, Testing performance of the RNN models on the DMS task. RNNs successfully trained either without noise (0 noise channels; n = 33) or with 10 noise channels (n = 50) were tested on the DMS task in which both internal noise and noisy input signals were introduced. We also varied the delay duration of these testing trials to range from 250 ms, 750 ms, and 1250 ms. For each testing condition, average accuracy of the trained RNN models is shown. Across all conditions, RNNs trained with no noise had lower accuracy than those trained with 10 noise channels (all *P s* < 0.01, two-sided Wilcoxon rank-sum test). Boxplot: central lines, median; bottom and top edges, lower and upper quartiles; whiskers, 1.5 *⇥* interquartile range; outliers are not plotted.

### Noise improves learning and enhances network resilience on working memory tasks

In order to study the effects of noise on the dynamics of the firing-rate RNNs and their performance on the DMS task, we introduced noise in the form of random Gaussian currents injected into the units during the training process (Fig. 1c; see *Methods*). For each noise level (C; see *Methods*), we trained 50 RNNs to perform the DMS task with a delay interval of 250 ms. Specifically, there were four stimulus conditions (s *2* {(+1, +1), (+1, *-*1), (*-*1, +1), (*-*1, *-*1)*}*). For the matched cases (stimulus condition 1 and 4), the model had to generate an output signal approaching +1. For stimulus condition 2 and 3 where the signs of the two sequential stimuli were opposite, the model had to produce an output signal approaching -1. As shown in Fig. 1d, the training success rate for the baseline model (i.e., no internal noise; C = 0) was 66% (33 out of 50 RNNs were trained within the first 20,000 trials). As the number of the noise channels (C) increased, the training success rate also increased (Fig. 1d). When C = 10, all 50 RNNs were successfully trained to perform the task (dark green in Fig. 1d). As shown in Extended Data Fig. 1, the internal noise with high variance (C = 1, 2, 4) did not provide any improvement in the training success rate compared to the RNNs trained without any noise (C = 0). Adding more than 10 noise channels (i.e., C = 20) did not result in additional improvement in the success rate. For the networks successfully trained, we did not see any significant difference in the number of training trials/epochs required among the four different noise conditions (Fig. 1e). We observed a similar trend for a DMS task involving two delay intervals (see *Methods*; Extended Data Fig. 2a-c).

As shown in Fig. 1d and Fig. 1e, the noise condition of C = 10 yielded the highest training efficiency. Importantly, the RNNs trained with this optimal noise structure were more robust to perturbations of internal dynamics compared to the RNNs trained without any injection of internal noise. Specifically, the RNNs trained with noise exhibited robust performance in the DMS task even when subjected to randomly generated internal noise introduced via randomly generated *w*^(noise)^, as opposed to the optimized *w*^(noise)^ used during training (Fig. 1f). In addition, the networks trained with noise demonstrated superior performance in the DMS task with a longer delay duration compared to the networks trained without noise (Fig. 1f). These results suggest that the injected noise facilitated contextualized sensory encoding and led to a more robust representation of the input stimuli.

To further investigate the impact of internal noise on the RNN dynamics, we applied the Potential of Heat-diffusion for Affinity-based Transition Embedding (PHATE; [19]) to the internal state trajectories of one example RNN realization from the baseline (C = 0) and noise (C = 10) conditions (see *Methods*). When only sensory noise (ξ in Equation (1)) was present, both models performed the DMS task with a delay of 250 ms equally well (Fig. 2a and Fig. 2b). However, applying PHATE to these two models revealed distinct differences in the dynamics and representations of the four stimulus conditions (Fig. 2c and Fig. 2d). In the RNN trained without noise, the neural representations of distinct stimulus conditions were found to intermingle in the lower-dimensional embedding space (Fig. 2c). However, in the RNN trained with noise (Fig. 2d), the dynamical structures corresponding to the four conditions were clearly demarcated, indicating a more distinct representation of the stimuli. Notably, these neural trajectories exhibited meaningful and informative bifurcations that were driven by the temporal structure of the DMS task (as indicated by the black arrows in Fig. 2d). Specifically, the first bifurcation occurred after presentation of the first stimulus (at 250 ms), followed by a second bifurcation at the onset of the second stimulus (at 750 ms). These distinct bifurcations observed in the trajectories over time highlight the role of internal noise in facilitating contextualized sensory encoding and working memory computation.

**Fig. 2:**
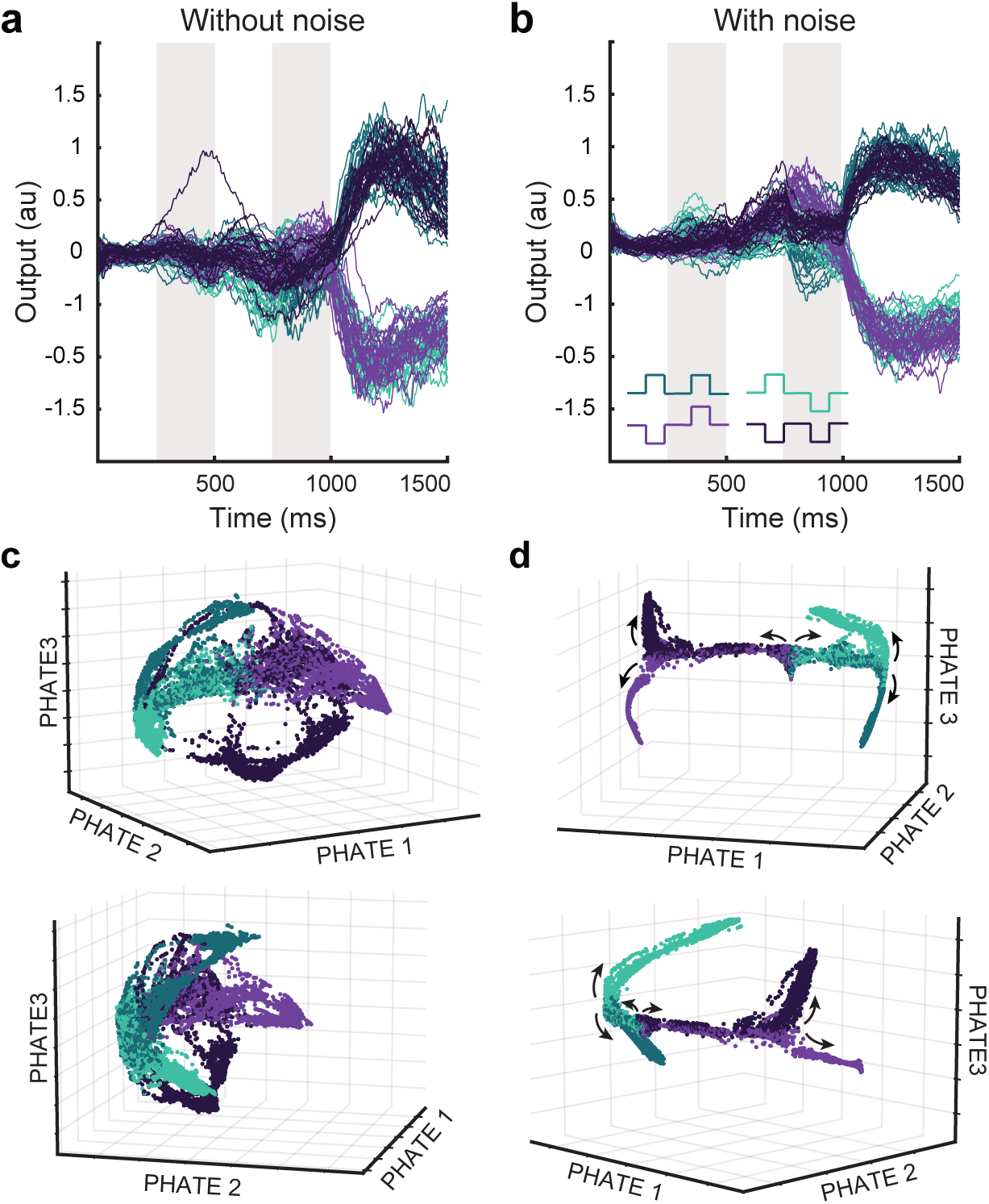
Neural representations of each stimulus condition on the DMS task. **a**, Network output of a sample RNN model successfully trained without noise to perform the DMS task with a delay of 250 ms. For match cases (s *2* {(+1, +1), (*-*1, *-*1)}), the network accurately generated an output signal approaching +1 (dark green and dark purple tracings). For non-match cases (s *2* {(+1, *-*1), (*-*1, +1)}), the network produced an output signal approaching -1 (light green and light purple). **b**, Network output of a sample RNN model trained with noise to perform the same DMS task as **a**. A schematic of the four stimulus conditions used in the DMS task shown in the bottom left corner. **c**, PHATE-embedding of the network activity (from the onset of the first stimulus window) from the RNN shown in **a**. **d**, PHATE-embedding derived from the network activity of the RNN trained with noise (same network as **b**). Black arrows indicate temporal progression of the PHATE trajectories over the trial duration.

### Noise modulates cell-type specific dynamics underlying working memory computation

Next, we investigated how the noise facilitated stable maintenance of stimulus information by examining the optimized model parameters. Given the previous studies highlighting the importance of inhibitory connections for information maintenance [9, 11–13], we hypothesized that the internal noise enhances working memory dynamics by selectively modulating inhibitory signaling. To test this, we first compared the inhibitory recurrent connection weights of the RNNs across different noise conditions (C = 0, 1, 5, 10). We did not observe any significant differences in the inhibitory weights (Extended Data Fig. 3). Similarly, the excitatory recurrent weights were also comparable across the noise conditions (Extended Data Fig. 3).

As we did not observe any noticeable changes in the recurrent weight structure induced by the noise, we next analyzed the distribution of the optimized synaptic decay time constants (τ). Interestingly, the synaptic decay constant distribution shifted toward the maximum value (125 ms; see *Method*) for the RNNs trained with noise (Fig. 3a). Separating the distribution of the inhibitory units from the excitatory units revealed that the change in the decay dynamics was mainly attributable to the shift in the inhibitory synaptic decay dynamics (Fig. 3b and Fig. 3c). In addition, the extent of the shift was correlated with the number of the noise channels (C): as C increased, the inhibitory synaptic signals decayed slower (Fig. 3c). Using noise with a higher variance also resulted in an increase in synaptic decay constants for both excitatory and inhibitory groups (Extended Data Fig. 4). These findings are in line with recent modeling studies that emphasized the importance of slow inhibitory dynamics in maintaining information [13].

**Fig. 3:**
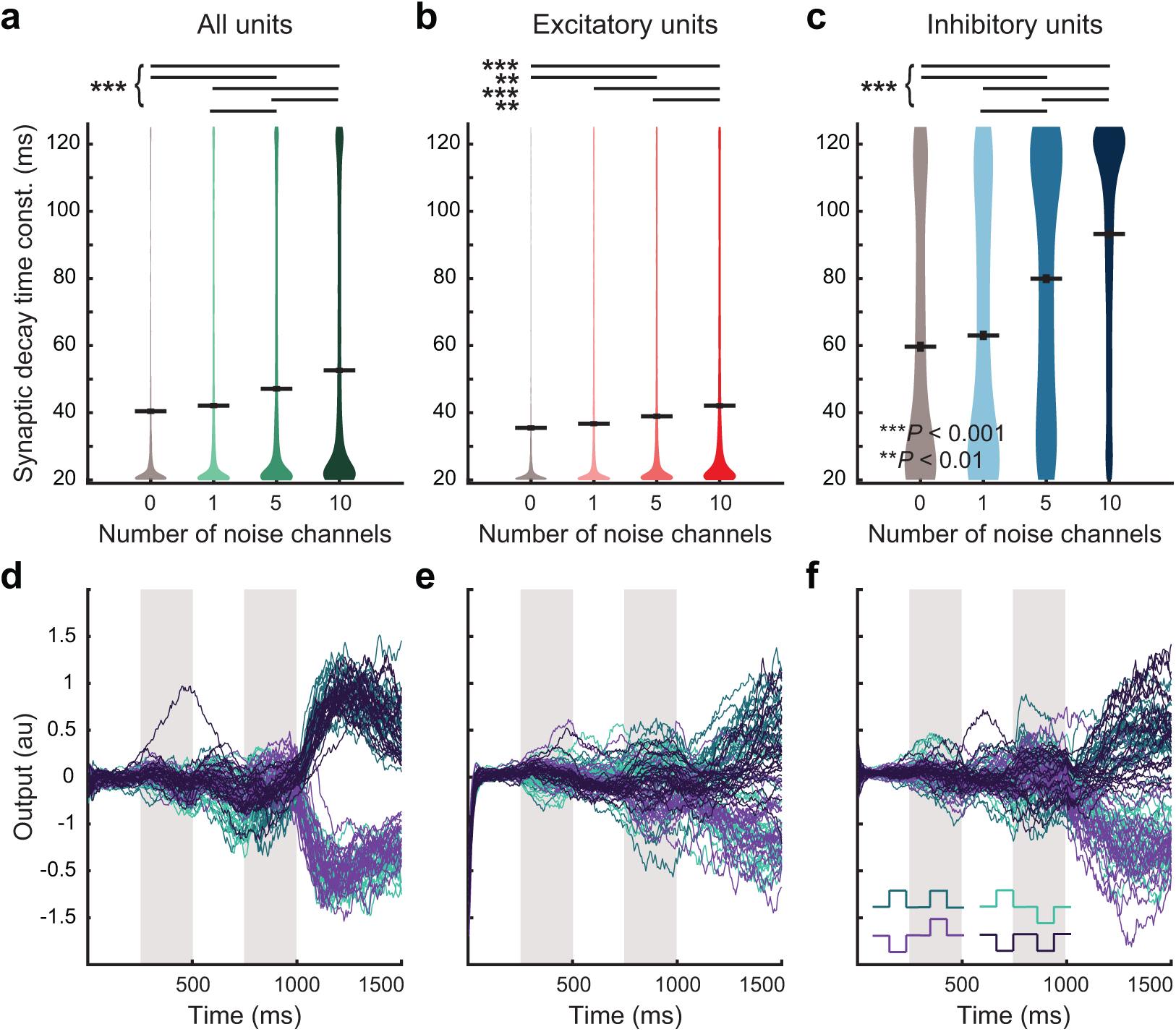
Influence of noise on cell-type specific temporal dynamics. Comparison of synaptic decay time constants of RNN models trained on the DMS task with varying amount of noise. **a**, For each noise condition, synaptic decay time constants of successfully trained models are reported for all units (n = 33, 40, 46, 50 for the noise conditions of 0, 1, 5, and 10 channels, respectively). Overall, injection of random noise during training increased synaptic decay time constants averaged across all units in the networks (*P s* < 0.001, *H* = 113.8; Kruskal-Wallis test with Dunn’s post hoc test). **b**, Comparison of synaptic decay time constants for excitatory units of the trained RNN models (*P s* < 0.01, *H* = 52.5; Kruskal-Wallis test with Dunn’s post hoc test). **c**, Comparison of synaptic decay time constants for inhibitory units of the trained RNN models (*P s* < 0.001, *H* = 120.3; Kruskal-Wallis test with Dunn’s post hoc test). Gray horizontal lines, mean. **d**, Network output of a sample RNN model trained without noise (same network as Fig. 2**a** and Fig. 2**c**). Re-plotted here for easier comparison. **e**, Network output of the same RNN model shown in **d** when synaptic decay time constants of all units were set to 125 ms (maximal τ; see *Methods*). The model failed to maintain memory and generate correct responses. **f**, Network output of the same RNN model shown in **d** when synaptic decay time constants of the inhibitory units were set to 125 ms (maximal τ; see *Methods*).

Since the RNNs trained with noise showed an increase in the inhibitory synaptic decay time-constant, we explored whether increasing the inhibitory τ would enhance the robustness of RNNs trained without noise. To test this hypothesis, we used the example RNN trained without noise (same network as the one shown in Fig. 2a). The network’s performance on the DMS task with a delay of 250 ms (Fig. 2a) is shown again in Fig. 3d for comparison. When τ for all the units in the network were increased to the maximum value (i.e., 125 ms), the network’s performance significantly decreased (Fig. 3e). We also observed that increasing the inhibitory τ to 125 ms, while keeping the excitatory τ at its original value, impaired the task performance (Fig. 3f). Together, these findings underscore the importance of incorporating internal noise during training to shape learned dynamics and enhance the network’s capacity to robustly perform working memory computations.

### Noise pushes model neurons with slow synaptic dynamics toward the edge of instability

Given that artificially increasing the inhibitory synaptic time constants in the RNNs trained without noise did not lead to improved memory maintenance (Fig. 3f), we next focused on understanding the role of slow inhibitory signaling in the networks trained with noise. For an RNN to perform well on the DMS task, it is plausible that RNN persistently maintains information during the delay window. This condition can be achieved when each unit in the network maintains relatively stable synaptic current activity throughout the delay window, i.e., *x*(*t*) *≈ x*^*^ at a given time point t during the delay period, where *x^*^* is the delay period steady state. For both models (RNNs trained without and with noise), the synaptic current activity during the delay period exhibited stability (Extended Data Fig. 5). We then performed the linear stability analysis around *x^*^*, revealing the role of slow inhibitory signaling as follows.

For each first stimulus condition, *s*_1_ ∈ {-1, +1}, we studied the impact of a small instantaneous perturbation around the stimulus-specific delay period steady state (*x*^*^_*s_1_*_). In the absence of an input stimulus and noise, the synaptic current activities evolve according to (modified from Equation (1)):

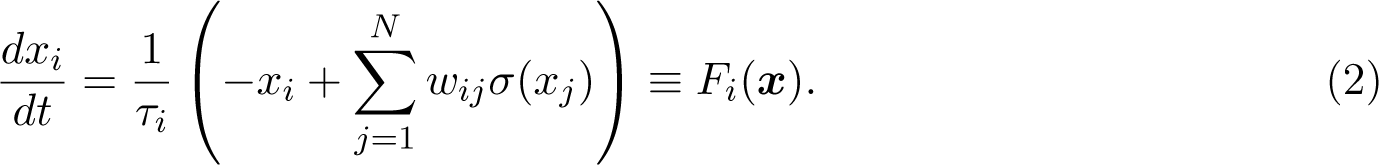

Perturbing *x^*^*_*s*_1__ by δ*x_s_*_1_ would lead to

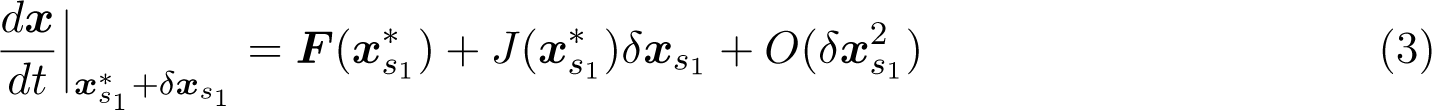

where *J*(*x*^*^_*s_1_*_) is the Jacobian matrix (see *Methods*). Since *F*(*x*^*^_*s_1_*_) *≈* 0, the perturbed dynamics *s*_1_ *s*_1_ (Equation (3)) can be re-written as

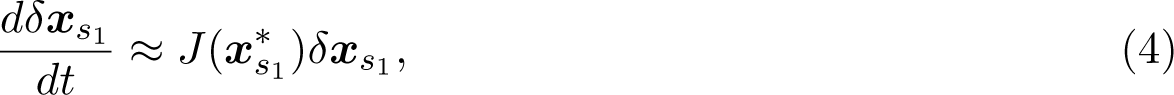

with the Jacobian matrix written explicitly as

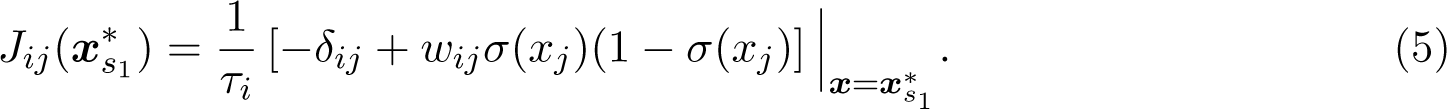

Performing spectral decomposition on *J* and calculating the eigenvalues (.X) of the example RNN models employed in Fig. 2 revealed that all eigenvalues of *J* exhibited negative real parts, indicating that the steady states (*x*^*^_*s_1_*_) are indeed stable against mild instantaneous perturbations (Fig. 4a–d; see *Methods*). Interestingly, the RNN model trained with noise contained more slowly relaxing modes with oscillatory behaviors compared to the network trained without noise (i.e., eigenvalues with non-zero imaginary components shifted toward zero along the real axis in Fig. 4c and Fig. 4d). Furthermore, these modes characterized by slow relaxation dynamics were found to exhibit de-localization, as evidenced by their low Inverse Participation Ratio (IPR) values (greener dots in Fig. 4c and Fig. 4d, and comparison of the average IPR values between the two RNNs shown in Fig. 4e; see *Methods*). Specifically, a larger IPR indicates a more localized perturbation that affects a smaller number of units, while a smaller IPR corresponds to a more delocalized perturbation affecting a larger number of units. In other words, RNNs trained with noise are more robust compared to the RNNs trained without noise, as they require sustained perturbations to a larger number of units for the steady state to be destabilized.

**Fig. 4:**
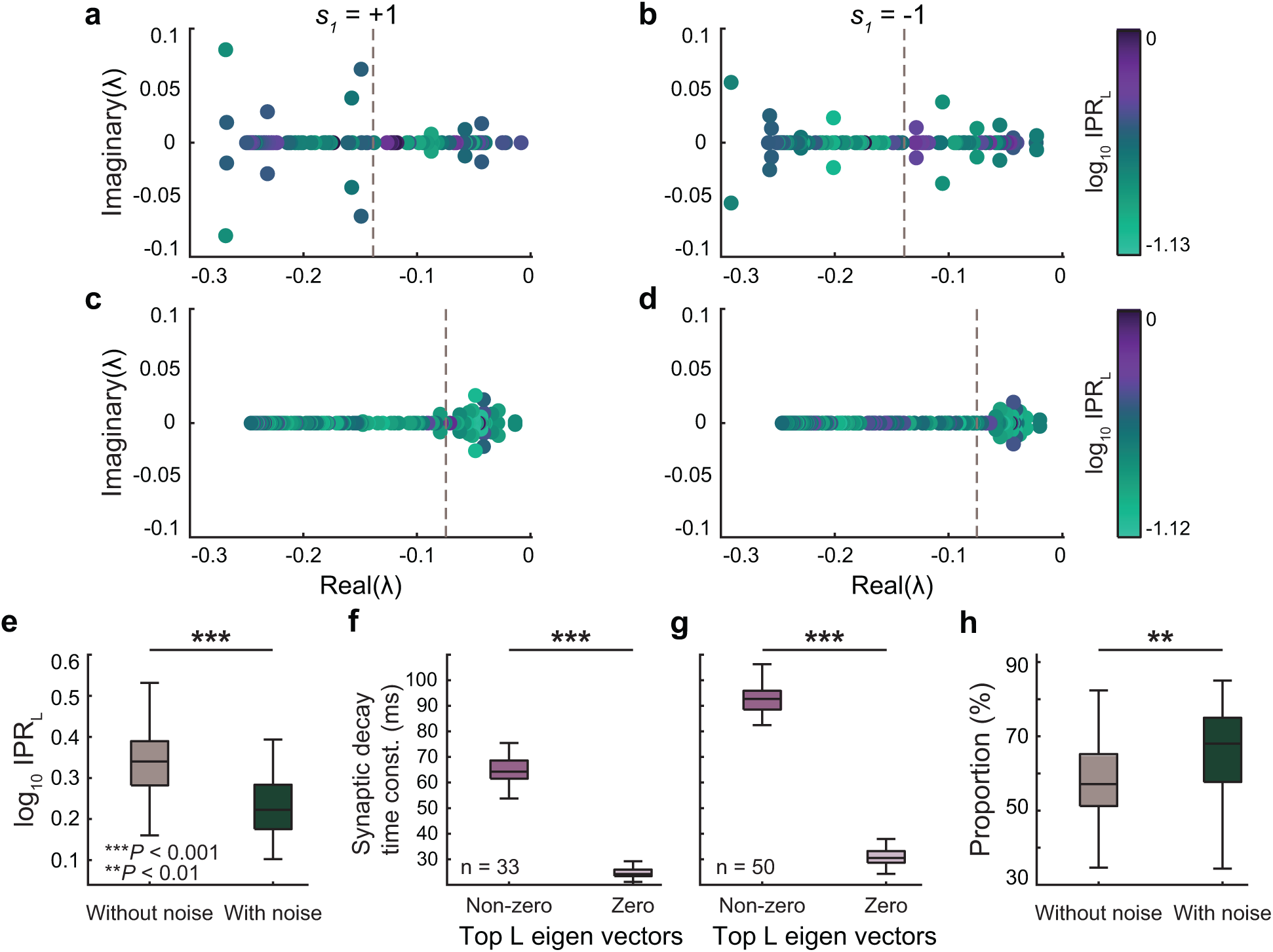
Spectral characteristics of network stability during the delay period. Spectra of the Jacobian (J) extracted from the network activity during the delay window. **a** and **b**, Spectra of a sample RNN model trained without noise (same RNN as Fig. 2**a**) during the delay period following the first stimulus presentation (*s*_1_ *ϵ* {+1, *-*1}). **c** and **d**, spectra of a sample RNN model trained with noise (C = 10; same network as Fig. 2**b**) during the delay period following the first stimulus presentation (*s*_1_ ϵ {+1, *-*1}). For both conditions, we observed stable steady states *x^*^* as evident from the real parts of all the eigenvalues being negative. For the RNN trained with noise, the eigenvalues with non-zero imaginary parts shifted to the right (toward zero along the real axis) and were associated with lower Inverse Participation Ratio (IPR) values (**c** and **d**). Vertical dashed lines represent the cutoff for the top 50 real eigenvalues. **e**, Average IPR values from the RNN trained without noise were significantly higher (i.e., more localized) than those from the model trained with noise. **f**, Average synaptic decay time constants of the dominant (non-zero elements in the top 50 eigenvectors) and non-dominant (zero elements in the top 50 eigenvectors) units from all the RNNs trained without noise. **g**, Average synaptic decay time constants of the dominant and non-dominant units from all the RNNs trained with noise. **h**, Proportion of inhibitory units among the dominant units in the RNNs trained with noise was significantly higher compared to the RNNs trained without noise. Boxplot: central lines, median; bottom and top edges, lower and upper quartiles; whiskers, 1.5 *⇥* interquartile range; outliers are not plotted. Two-sided Wilcoxon rank-sum tests were performed.

In order to further characterize the slow relaxation modes observed in the RNN trained with noise, we first identified the units involved in the left eigenvectors corresponding to the top 50 eigenvalues (i.e., 50 least negative eigenvalues) for each RNN model (see *Methods*). We categorize the units with non-zero amplitudes in the top 50 eigenvectors as dominant units (perturbation on these units could more dominantly influence the RNN to destabilize), while the units with zero amplitudes are referred to as non-dominant units. Notably, in both RNN models (trained without and with noise), the dominant units were associated with significantly larger synaptic decay time constants compared to the non-dominant units (Fig. 4f and 4g). Furthermore, the synaptic decay dynamics of the dominant units in the RNNs trained with noise were significantly slower than the dynamics of the dominant units in the networks trained without noise (P < 0.001, two-sided Wilcoxon rank-sum test). Interestingly, the top 50 left eigenmodes from the RNNs trained with noise contained a significantly larger number of inhibitory units than the top eigenmodes from the networks trained without internal noise (Fig. 4h).

These findings suggest that injection of the internal noise during training resulted in an increased proportion of units exhibiting slower synaptic dynamics (i.e., dominant units). In addition, this noise-induced effect pushed the top left eigenmodes composed of these units closer to the edge of instability (critical boundary between stable and unstable behavior).

To investigate the impact of these factors on working memory, our analysis focused on the sustained maintenance of *s*_1_ = +1 during a long delay period (1250 ms) in both models. Since the networks consisted of units that were selective to each of the first stimulus conditions (*s*_1_ ϵ{+1, *-*1}), the successful maintenance of *s*_1_ = +1 during the delay period relied on two key conditions: persistent excitation of the units tuned to *s*_1_ = +1 and persistent inhibition of the units tuned to *s*_1_ = *-*1. As shown in Fig. 5a, the average normalized firing rate timecourses (normalized by subtracting the average baseline activity; see *Methods*) of the dominant units preferring *s*_1_ = +1 in the top 50 left eigenmodes of the RNNs trained without noise demonstrated higher firing rates during the stimulus presentation of +1 compared to the non-dominant units selective for *s*_1_= +1. Throughout the delay period, the average normalized firing rate activity of the dominant units exhibited a rapid decay (Fig. 5a). Repeating the above analysis on the RNNs trained with noise revealed that the dominant units maintained *s*_1_ = +1 at a significantly higher rate throughout the delay window than the dominant units from the networks trained without noise (Fig. 5b; *P* < 0.001, two-sided Wilcoxon rank-sum test), consistent with the slow synaptic dynamics seen in Fig. 4.

**Fig. 5:**
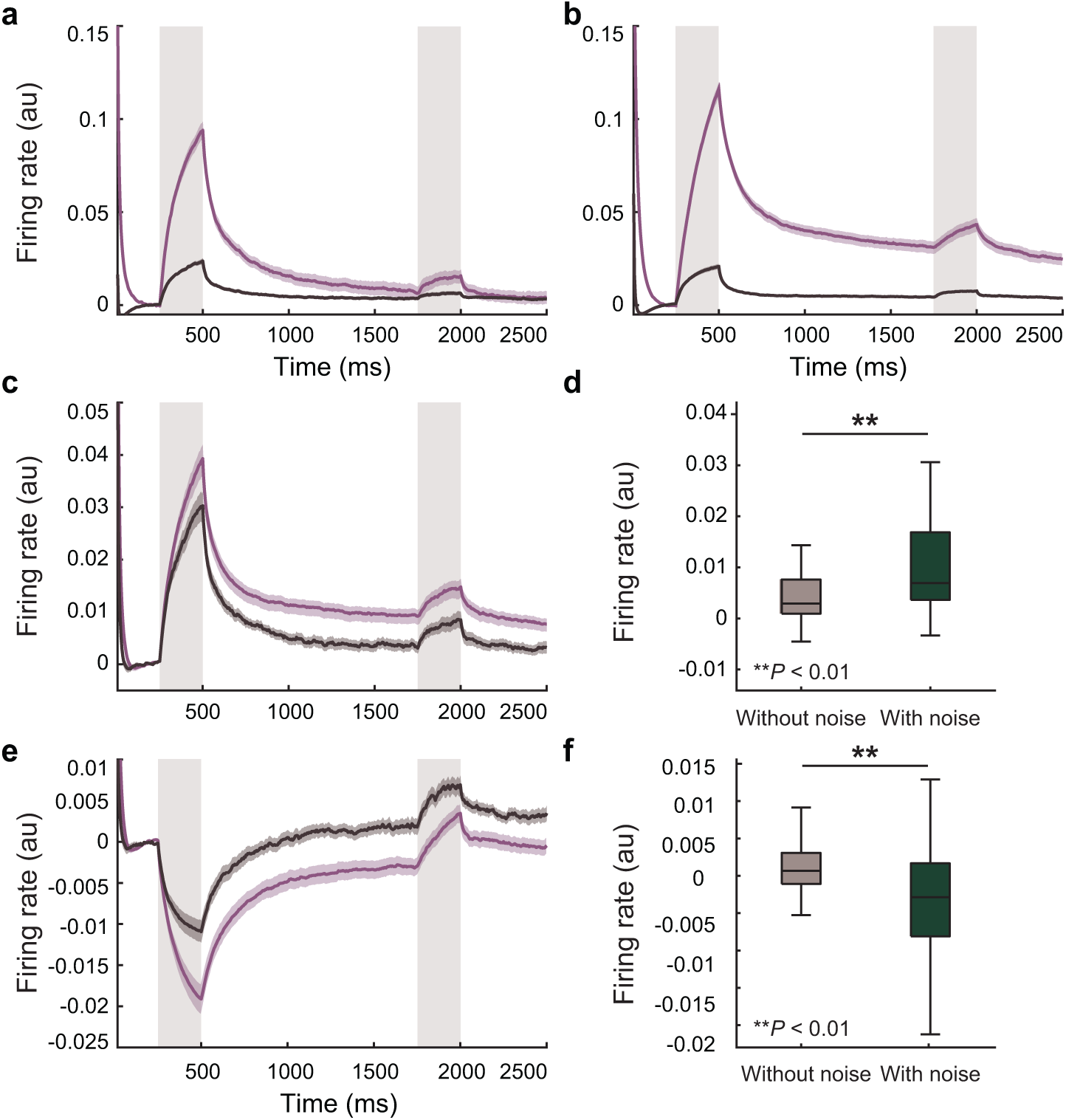
Persistent activity of dominant units from RNNs trained with noise. **a**, Average normalized firing rate timecourses of the dominant units selective to +1 (purple) and non-dominant units preferring +1 (dark gray) in the top 50 left eigenmodes of the RNNs trained without noise. The DMS task was modified to have a delay duration of 1250 ms, and 50 trials with the first stimulus of +1 were used to extract the timecourses. **b**, Average normalized firing rate timecourses of the dominant units selective to +1 (purple) and non-dominant units preferring +1 (dark gray) in the top 50 left eigenmodes of the RNNs trained with noise. **c**, Average normalized firing rate timecourses of the units preferring +1 in the corresponding right eigenvectors from the RNNs trained without noise (dark gray) and with noise (purple). **d**, Average normalized firing rate of the units shown in **c** during the late delay period (last 750 ms) was significantly higher for the RNNs trained with noise compared to the networks trained without noise. **e**, Average normalized firing rate timecourses of the units preferring -1 in the corresponding right eigenvectors from the RNNs trained without noise (dark gray) and with noise (purple). **f**, Average normalized firing rate of the units shown in **e** during the late delay period (last 750 ms) was significantly lower for the RNNs trained with noise compared to the networks trained without noise. Mean *±* standard error shown. Boxplot: central lines, median; bottom and top edges, lower and upper quartiles; whiskers, 1.5 *⇥* interquartile range; outliers are not plotted. Two-sided Wilcoxon rank-sum tests were performed.

Next, we directed our attention to the units in the corresponding right eigenmodes to investigate the impact of the slow dynamics observed in the top left eigenvectors on the dynamics of the network response (see *Methods*). We hypothesized that the slow decay and persistent activity observed in the units of the top left eigenmodes would confer similar properties to the units in the corresponding right eigenvectors during the delay period. Additionally, we posited that the units tuned for +1 and -1 in the right eigenmodes would exhibit persistent excitation and inhibition, respectively. As shown in Fig. 5c and Fig. 5d, the units tuned for +1 in the right eigenmodes of the RNNs trained with noise demonstrated significantly higher activity during the delay period compared to the +1-preferring units in the right eigenmodes of the networks trained without noise. Furthermore, the units tuned for -1 in the right eigenmodes of the RNNs trained with noise exhibited significant suppression throughout the delay period compared to the units tuned for -1 in the right eigenmodes of the networks trained without noise (Fig. 5e and Fig. 5f). These results suggest that the networks trained with noise exhibit greater robustness to perturbations compared to the RNNs trained without noise, and the noise-induced increase in synaptic decay time constants of the inhibitory units near the edge of instability facilitated maintenance of stimulus-specific information for an extended duration.

### Robustness and increased efficiency due to internal noise are specific to working memory computations

Finally, we asked if the modulatory effects of noise during training were specific to working memory dynamics. To address this question, we devised two cognitive tasks that do not require maintenance of sensory information over time, namely two-alternative forced choice (AFC) task and context-dependent sensory integration (CTX) task (see *Methods*). In the AFC task (Fig. 6a), the RNN model had to generate an output signal that indicated whether a target sensory signal was present. The CTX task is a more challenging variant of the AFC task, where the model was trained to produce an output that corresponded to one of the two input modalities as determined by a context signal [14] (Fig. 6b). As these task paradigms do not involve any delay interval, the model only requires minimal information maintenance, if any, to perform well on these tasks.

**Fig. 6:**
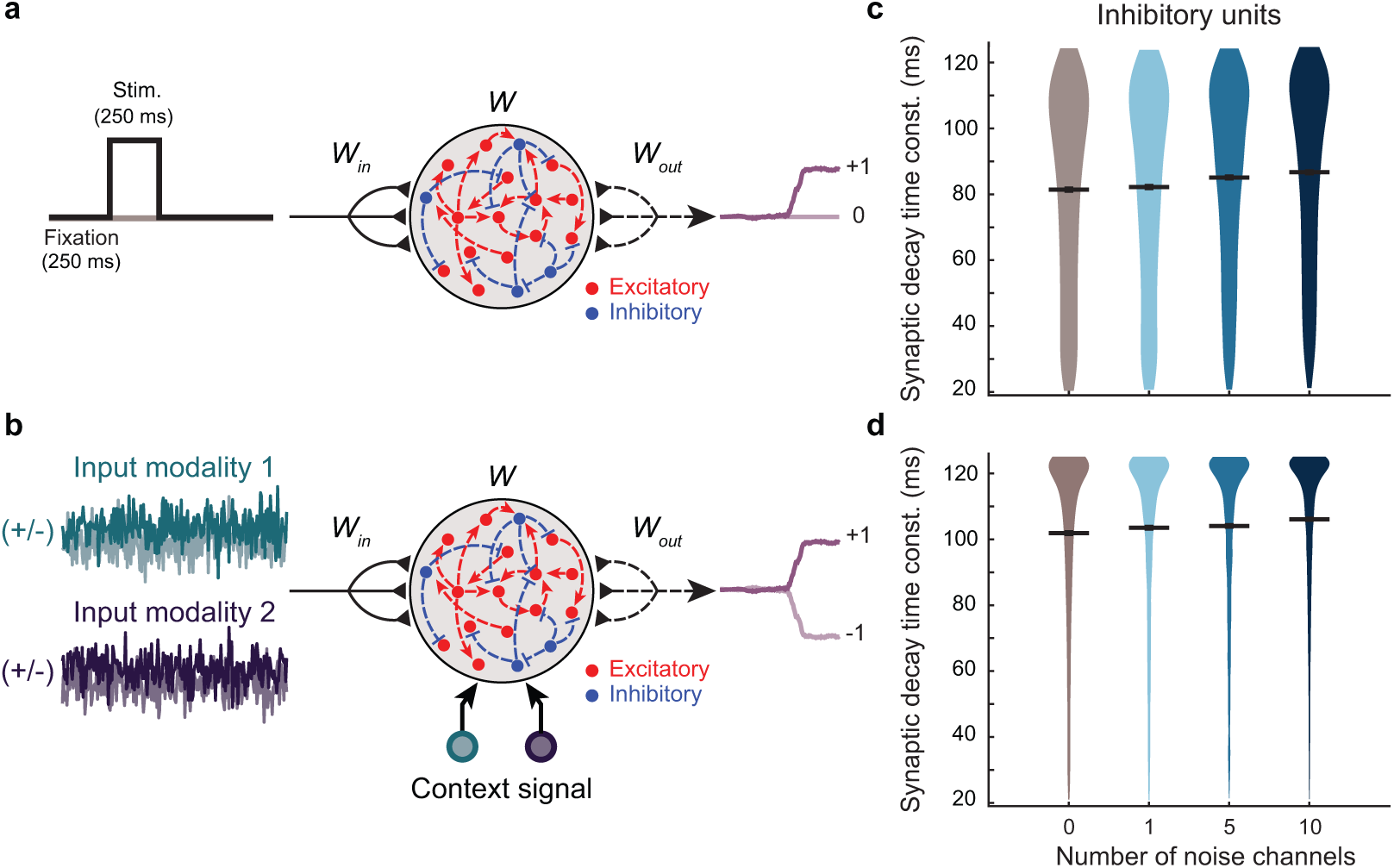
Network functional motifs underlying working memory-independent computation. Schematics diagrams illustrating working memory-independent tasks and the corresponding network dynamics of the RNN models successfully trained on these tasks. **a**, Two-alternative forced choice (AFC) task, in which the RNN modes were trained to produce an output indicating the presence of a brief input pulse. **b**, Context-dependent sensory integration (CTX) task, where the RNN models were trained to generate an output based on the identity of a sensory stimulus whose relevance was determined by an explicit context cue. **c**, For the inhibitory units from the RNNs trained on the AFC task, synaptic decay time constants were similar across all noise conditions. **d**, Across all the noise conditions, the inhibitory units from the networks trained on the sensory integration task exhibited similar synaptic decay time constants. Gray horizontal lines, mean.

Our findings demonstrated that the RNN models were able to perform these non-working memory tasks well without any noise, and that adding noise during training did not further improve training efficiency. In fact, it took longer for models to reach successful training criteria when noise was added during training for both sensory integration and context-dependent sensory integration tasks (*P s* < 0.001 for both tasks). To investigate if noise modulated the temporal dynamics on these tasks, we analyzed synaptic decay time constants of all the units as well as separately for excitatory and inhibitory units. Our results revealed no difference in the synaptic decay dynamics in the inhibitory units from the models that trained without noise and those trained with noise (Fig. 6c and d). These findings suggest that the slow synaptic decay dynamics induced by noise are specific to working memory functioning where robust information maintenance is needed to ensure successful performance.

## Discussion

In this study, we demonstrated that introducing random noise into firing-rate RNNs allowed the networks to achieve efficient and stable memory maintenance critical for performing working memory tasks. We also showed that the models trained with noise were able to generalize to sustain stimulus-related information longer than the delay period used during training. Further analyses uncovered that the introduction of noise led to the emergence of inhibitory units with slow synaptic decay dynamics, which were predominantly associated with dominant eigenmodes situated near the edge of instability. These eigenmodes were critical for maintaining information during the delay period of the working memory task. Specifically, the network should exhibit stability to prevent minor noisy perturbations from causing substantial alterations in its dynamics and compromising the information of the stimulus. However, the network should not be overly stable, as this would result in rapid decay of the information associated with the stimulus, as the network quickly returns to its stable configuration. Hence, these eigenmodes crucial for robust memory maintenance emerge near the edge of instability. In addition, these effects were specific to the models trained to perform working memory task, suggesting that noise-induced changes were specific to working memory.

Our findings are closely related to the previous studies that reported the benefits of random neural noise ubiquitous in the cortex in memory recall and associative learning [20, 21]. For example, recent experiments showed that a high level of noise and randomness in the olfactory system (i.e., random and seemingly unstructured networks in the piriform cortex) allows for not only flexible encoding of sensory information but also maintenance of the encoded information [4, 22–24]. Consistent with this line of work, the injected noise in our RNN models during the training helped stabilize the encoding of sensory space and thus enhanced learning efficiency. Taken together, our study provides an easy-to-use framework for understanding how internal noise influences information maintenance and learning dynamics when performing working memory cognitive tasks.

One limitation of the present study is the lack of comparisons with RNNs trained with learning algorithms that are not based on gradient-descent optimization. One such algorithm is First-Order Reduced and Controlled Error (FORCE) learning which has been employed to train rate and spiking RNNs [25, 26]. Due to the nature of the method, it is currently not possible to train the synaptic decay time constant term using FORCE training, making the comparison with our models difficult. Reinforcement learning is another learning algorithm that can be employed to train biologically realistic RNNs [27].

Even though we showed that increasing the number of noise channels could lead to heterogeneous synaptic decay time constants, it is unclear why only inhibitory synaptic decay constants undergo significant changes for working memory tasks. Future work will focus on better understanding the theoretical and computational basis for the emergence of slow inhibitory synaptic dynamics.

By interpreting the concept of noise in machine learning within the context of biology, the present study proposes a general framework that bridges recent advances in machine intelligence with empirical findings in neuroscience. Our approach includes introducing internal noise into a biologically realistic artificial neural network model during training to simulate cortical noise and systematically evaluating its effects on model dynamics and performance under different testing conditions. Elucidating the computational underpinnings of how cortical noise modulates cognitive functions will help us better understand how such processes are disrupted in neuropsychiatric conditions such as schizophrenia and autism spectrum disorder. Finally, our framework has the potential to shed light on the fundamental mechanisms that may give rise to the therapeutic effects of deep brain stimulation (DBS), a neuromodulation technique that entails the targeted delivery of electrical stimulation to specific brain regions.

## Methods

### Continuous-rate recurrent neural network (RNN) model

We constructed our biologically realistic RNN model based on Equation (1). All the units in the network are governed by the following equations:

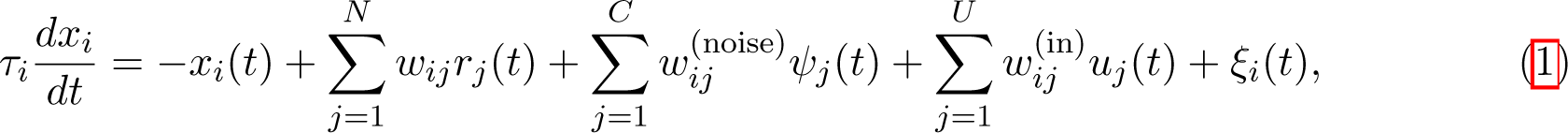

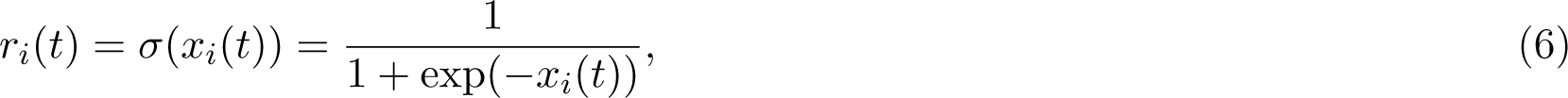

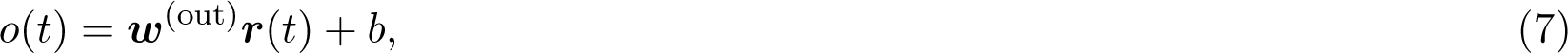

where τ*_i_* is the synaptic decay time constant of unit *i*, *x_i_* is the synaptic current variable of unit *i, w_ij_* is the synaptic weight from unit *j* to unit *i*, and *r_i_* is the firing rate estimate of unit *i* (estimated by using the sigmoid transfer function in Equation (6)). Each model contains 200 units. To adhere to previous empirical observations regarding the proportion of excitatory and inhibitory units in the brain, we constructed each RNN with a composition of 80% excitatory and 20% inhibitory units (i.e., E-I ratio of 80/20; [28–30]). The model receives time-varying input composed of U channels of signals over T time steps (*u* ϵR*^U⇥T^*) via the input weight matrix, *w*^(in)^ ϵR*^N⇥U^*. For the DMS task, u contained two streams of input signals (i.e., *U* = 2). The network also receives random noise via *w*^noise^ ϵR*^N⇥C^* where C is the number of independent noise signals in ϵR*^C⇥T^*. Each signal in was independently drawn from the standard Gaussian distribution with zero mean and unit variance. We considered C ϵ{0, 1, 5, 10} in this study. The sensory noise (ξ ϵR*^N⇥T^* ; uncorrelated in time) was generated from a Gaussian distribution with zero mean and a variance of 0.01. The output (o) of the network was computed as a weighted average of the activities of the units via the readout weights (*w*^(out)^) and the constant bias term (b).

The dynamics were discretized using the first-order Euler approximation method and with the step size (i6.t) of 5 ms:

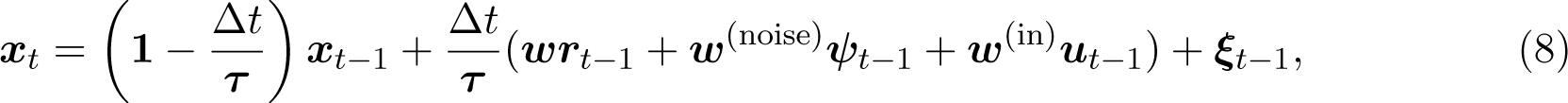

where 1/τ denotes a diagonal matrix whose i^th^ diagonal element is 1/τ*_i_*. The network was trained using backpropagation through time (BPTT). The trainable parameters of the model included w, *w*^(noise)^, τ, w^(out)^, and b. To further impose biological constraints, we incorporated Dale’s principle (separate populations for excitatory and inhibitory units) using methods similar to those implemented in previous studies [15, 31].

Instead of fixing the synaptic decay constant (τ) to a fixed value for all the units, we optimized the parameter for each unit using a similar algorithm similar to the method described in Kim et al. [31]. The parameter was trained to range from 20 ms to 125 ms to model heterogeneous synaptic dynamics of different receptors in the cortex [32, 33]. We initialized the synaptic decay time constant parameter (τ) using τ*_i_* = o(*N* (0, 1))τ*_step_* + τ*_min_*, where o(*·*) is the sigmoid function and *N* (0, 1) refers to the standard normal distribution. τ*_min_* = 20 ms and τ*_step_* = 105 ms were used to constrain the parameter to range from 20 ms to 125 ms. The gradient of the cost function with respect to the synaptic decay term is derived in *Supplementary Information*.

The schematic diagram of the model is shown in Fig. 1c. All the models were implemented with TensorFlow 1.10.0 and trained on NVIDIA GPUs (Quadro P4000 and Quadro RTX 4000).

### Delay match-to-sample (DMS) task

Two match-to-sample (DMS) tasks were used to train our RNN model and assess how the noise influenced the robustness of memory maintenance in the network. Both tasks involved two sequential stimuli (each lasting 250 ms) separated by a delay interval of 250 ms. The first stimulus was presented after a fixation period of 250 ms. During the stimulus window, the input signal (u) was set to either -1 or +1 (Fig. 1a). If the signs of the two sequential stimuli matched (i.e., stimulus condition 1: *s* = (+1, +1); stimulus condition 4: *s* = (*-*1, *-*1); Fig. 3a), the model was trained to produce an output signal approaching +1. When the signs were opposite (i.e., stimulus condition 2: *s* = (+1, *-*1); stimulus condition 3: *s* = (*-*1, +1); Fig. 3a), the model had to produce an output signal approaching -1. For the first task, the model had to respond immediately after the second stimulus (Fig. 1c). A second delay period of 250 ms was added after the second stimulus for the second task (Extended Data Fig. 2a). Due to the two delay periods, the second DMS task is considered a more challenging working memory task than the first task. The primary focus of the present study is the one-delay DMS task, and all the DMS findings presented in the main text are exclusively derived from this specific paradigm. The results for the two-delay DMS task are shown in Extended Data Fig. 2.

### Training protocol

Our model training was deemed successful if the following two criteria were satisfied within the first 20,000 epochs:

- Loss value (defined as the root mean squared error between the network output and target signals) < 7
- Task performance (defined as the average accuracy of the network output over 100 randomly generated testing trials) > 95%

If the network did not meet the criteria within the first 20,000 epochs, the training was terminated. For each task and each value of C ϵ{0, 1, 5, 10}, we trained 50 RNNs using the above strategy. We considered the RNNs trained with C =0 (i.e., without any noise) as the baseline model.

### Testing protocol

To evaluate the robustness and stability of the trained RNNs, we devised a series of testing conditions where different aspects of the one-delay DMS task (Fig. 1f) were systematically manipulated. During testing, internal noise and noisy input signals were introduced to the trained networks. For each successfully trained RNN, we generated *w*^(noise)^ and as identically distributed Gaussian random variables to deliver random noise during testing.

For the noisy input signal, white-noise signals (drawn from the standard normal distribution) were added to the sensory signals (u) to mimic stimulus-related noise. Additionally, we also varied the duration of the delay interval to range from 250 ms to 1250 ms (with a 500-ms increment) to assess the stability of memory maintenance (Fig. 1f).

### Working memory-independent tasks

In addition to the DMS tasks that require memory maintenance over time, we designed two additional cognitive tasks that do not involve working memory computation. By comparing the dynamics of the RNNs between the DMS tasks and these working memory-independent tasks, we were able to identify the specific network dynamics associated with working memory computation.

For the two-alternative forced choice (AFC) task, our RNN model was trained to produce an output signal approaching +1 when a stimulus was presented (250 ms in duration), following a fixation period of 250 ms. For a trial where a stimulus was not presented, the model had to maintain the output signal close to 0 (Fig. 6a). For the context-dependent sensory integration (CTX) task, the model received two streams of noisy stimulus signals (input modality 1 and input modality 2; (Fig. 6c) along with a constant-valued, context signal which informed the model which sensory input modality was relevant on each trial. A random Gaussian time series signal with zero mean and unit variance was used to simulate a noisy sensory input signal. Each time series signal was then shifted by a positive or negative constant offset value to encode sensory evidence towards either the positive or negative choice, respectively. The magnitude of the offset value determined the degree of evidence for the specific choice (positive/negative) represented in the relevant noisy input signal. The network had to generate an output signal approaching +1 or -1 in response to the cued input signal with a positive or negative mean, respectively. Thus, if the cued input signal was generated with a positive offset value, the network was expected to produce an output that approached +1 irrespective of the mean of the irrelevant input signal. For both the AFC and CTX tasks, the training termination criteria were similar to those used for the DMS (see *Training protocol*).

### Visualization of network dynamics

To visualize the neural dynamics of working memory computation as a function of injected internal noise during training, we employed the Potential of Heat-diffusion for Affinity-based Transition Embedding (PHATE) algorithm [19]. This dimensionality reduction technique is a manifold learning algorithm that enables faithful visualization of high-dimensional data while best preserving the global data structure. Two example RNN models successfully trained either without (C = 0) or with noise (C = 10) were presented with a simulation of 100 DMS test trials (25 from each of the four stimulus conditions). The delay interval was fixed at 250 ms, such that the temporal structure of the testing phase mirrored that of the training environment (see Fig. 1c).

We then used the resulting neural activity data from each model type during this testing phase as input data for PHATE in order to compute the low-dimensional embedding corresponding to the neural activity of the RNNs trained with and without noise. Specifically, for each of the RNNs trained under each noise condition (without or with noise), the diffusion operator matrix was first calculated using pairwise similarities among individual points in the input network activity time series (downsampled by a factor of 5). This matrix was raised to a power exponent to amplify the local structure while preserving the global structure of the input data. The resulting matrix was then used to generate the low-dimensional embedding that captures the neural dynamics of the input data.

To characterize potential topological patterns within the neural dynamics associated with each RNN, clustering was performed on this PHATE-generated embedding. Specifically, a K-means clustering algorithm was used to partition the data into distinct groups based on their spatial proximity in the low-dimensional space. For visualization purposes, a 3-dimensional PHATE embedding of a sample model from each noise condition (i.e., without noise and with noise; Fig. 2c and Fig. 2d) was plotted and colored by stimulus conditions. Black arrows were also included to indicate the temporal evolution of the neural trajectories over the trial duration. These embeddings provided insights into the temporal structure underlying working memory computation associated with the network dynamics that resulted from the incorporation of internal noise during training.

### Network stability analysis during the delay interval

To investigate the neural dynamics associated with memory maintenance, we employed linear stability analysis. Specifically, we performed this analysis on the synaptic currents of the RNNs successfully trained without or with noise during the delay period in the DMS task (i.e., from the offset of the first stimulus to the onset of the second stimulus (see Fig. 1c). Throughout this window, the network activities exhibited consistent steady-state patterns, as illustrated in Extended Data Fig. 5.

For each first stimulus condition *s*_1_ *2* {-1, +1}, we defined the steady-state synaptic current variable (*x*^*^_*s_1_*) by_ first averaging x*_s_* (*t*) across time within the delay window and then averaging across *s*_1_ 1 multiple trials (50 trials per each first stimulus condition). The impact of a small instantaneous perturbation around the delay period steady state *x^*^* on the synaptic current patterns is determined by the deterministic dynamics of Equation (1) in the absence of an input stimulus:

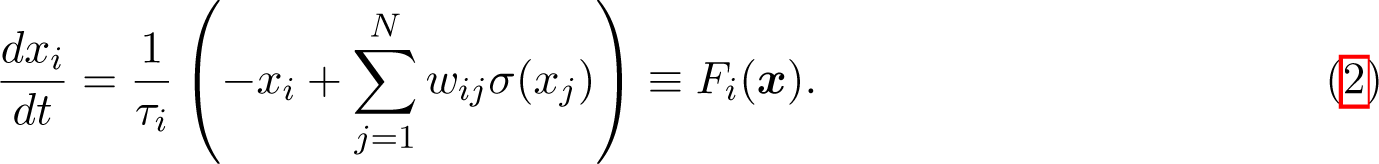

For a weak perturbation bx*_s_*_1_ around x, the linearized approximation of the perturbed dynamics 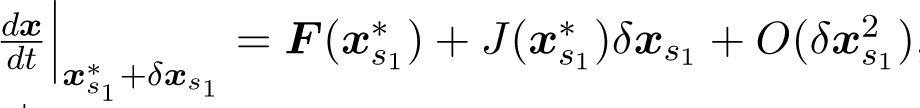, where *J*(*x*^*^_*s_1_*_) is the Jacobian matrix 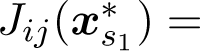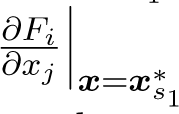. By the assumption of the steady state *x*^*^_*s_1_*_, which is also consistent with the numerical results, we have ***F*** (*x^*^_*s*1_*) *≈* **0**. Thus, the linearized dynamics of the perturbation bx*_s_*_1_ can be written as

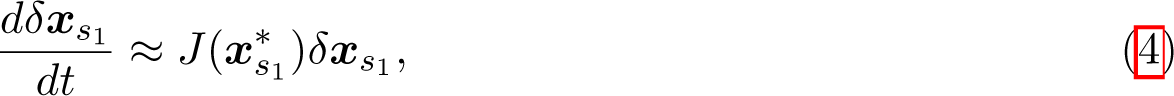

with the Jacobian matrix written explicitly as

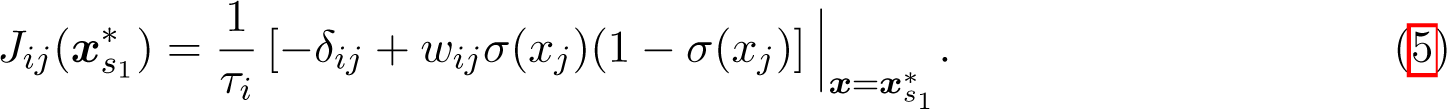

Network responses to weak perturbations around the steady states can now be systematically explored by the spectral analysis (eigenvalues and eigenvectors) of the Jacobian in Equation (5).

For brevity, we will add the subscript s only when the stimuli-specific statement is needed. Also, *J* will denote the Jacobian evaluated at the steady state of interest. In this notation, given the linearized perturbed dynamics of Equation (4), the initial perturbation *δx_0_* will evolve into the response at time t, bx(*t*), that can be studied via the spectral decomposition of *J* [34] as

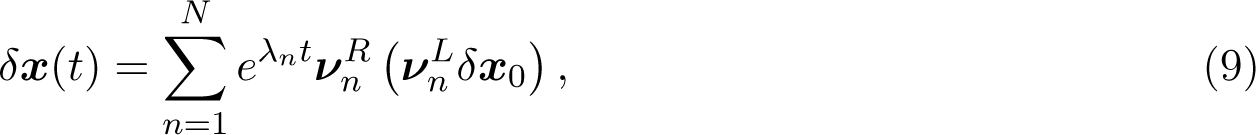

where *v^L^_n_* and *v^R^_n_* are, respectively, the left and the right eigenvector of *J* with the eigenvalue *λ_n_*. Notably, our trained RNNs exhibit highly asymmetric w such that the Jacobian Equation (5) is non-hermitian, leading to distinct left and right eigenvectors.

Equation (9) states that an initial perturbation *δx_0_* via *v^L^_n_* will contribute to a response *v^R^_n_*, such that the response will grow (decay) exponentially on the timescale of |1/Re (A*_n_*)| when Re (A*_n_*) > 0 (Re (A*_n_*) < 0).

Since the dominant responses to a perturbation depend on the overlap between the perturbation and the top-most left eigenvectors *v^L^_n_*δx_0_, the non-zero elements of the top-most left eigenvectors determine the spatial extent of perturbation required to significantly influence the system’s response. Along this line, the larger the number of non-zero elements in the top-most left eigenvectors, the larger the number of units that need to be perturbed to destabilize the steady states.

We employ the Inverse Participation Ratio (IPR), a measure commonly used in the study of localization phenomena in statistical physics [35], to reflect the number of units participating in the perturbation. The IPR provides valuable insights into the localization of perturbations by indicating the number of units involved in the perturbation process. In particular,

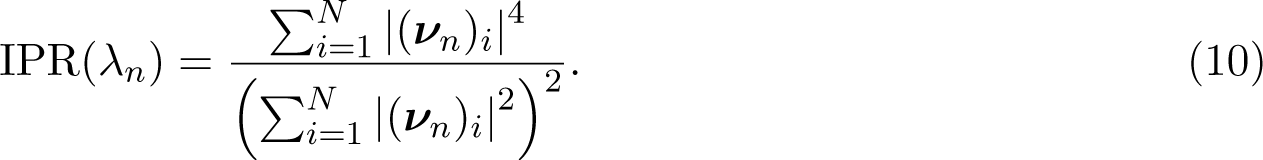

The IPR of the left and the right eigenvector will be denoted by IPR*_L_* and IPR*_R_* respectively, though we will focus on IPR*_L_* as we are interested in the size of the neural subpopulations participating in the perturbation. Note that the maximum and the minimum values of IPR*_L_* are attained at, respectively, 1 when only a single neuron is non-zero, and 1/N when all the units are uniformly activated. A larger or a smaller value of IPR*_L_* indicates that the perturbation is localized around a smaller number of units, or extended over a larger number of units, respectively.

### Stimulus selective units

To identify units selectively tuned for each of the first stimulus condition (*s*_1_ ϵ{+1, *-*1}), we first generated 50 trials for each stimulus condition and computed average firing rates (r) during the first stimulus presentation window. Next, we performed a one-sided Wilcoxon rank-sum test for each unit to determine its selectivity.

### Firing rate normalization

For the firing rate timecourses (Fig. 5), we normalized the trial-averaged firing rate of each unit by subtracting its corresponding baseline trial-averaged firing rate. The baseline activity was determined by considering the window preceding the onset of the first stimulus.

### Statistical analyses

All the RNNs trained in the present study were randomly initialized (with random seeds) before training. Throughout this study, we employed non-parametric statistical methods to assess statistically significant differences between groups. For comparing differences between two groups (e.g., the log_10_ IPR_L_ of RNNs trained with or without noise), we used two-sided Wilcoxon rank-sum or signed-rank test. For comparing more than two groups (e.g., the synaptic decay time constants associated with RNNs trained with varying degree of noise), we used Kruskal-Wallis test with Dunn’s post hoc test to correct for multiple comparisons.

## Acknowledgements

This work was supported by the Swartz Foundation (N.R.), the Kavli Institute for Brain and Mind (N.R.), the National Institute of Biomedical Imaging and Bioengineering R01EB026899-01 (T.J.S.), National Institute of Neurological Disorders and Stroke R01NS104368 (T.J.S.), Mission funding from the cooperative agreement under the United States Army Research Laboratory W911NF-17-S-0003-03 (N.R.), the funding from the National Research Council of Thailand #1187111 on the fiscal year 2023 (T.C.), and from Thailand Science Research and Innovation Fund Chulalongkorn University (IND66230005) (T.C.) The views and conclusions contained in this document are those of the authors and should not be interpreted as representing the official policies, either expressed or implied, of the DEVCOM Army Research Laboratory or the United States Government.

## Author information

**Department of Biomedical Engineering, Columbia University, New York, NY, USA**

Nuttida Rungratsameetaweemana

**Computational Neurobiology Laboratory, Salk Institute for Biological Studies, La Jolla, CA, USA**

Nuttida Rungratsameetaweemana, Robert Kim & Terrence J. Sejnowski

**Neurology Department, Cedars-Sinai Medical Center, Los Angeles, CA, USA** Robert Kim

**Chula Intelligent and Complex Systems, Department of Physics, Chulalongkorn University, Bangkok, Thailand**

Thiparat Chotibut

**Institute for Neural Computation, University of California San Diego, La Jolla, CA, USA**

Terrence J. Sejnowski

**Division of Biological Sciences, University of California San Diego, La Jolla, CA, USA**

Terrence J. Sejnowski

## Contributions

N.R. and R.K. conceived, designed, and performed the research; N.R., R.K., and T.C. analyzed data; N.R., R.K., T.C., and T.J.S. wrote the manuscript.

## Corresponding author

Corresponding authors: correspondence to Thiparat Chotibut or Terrence J. Sejnowski

## Code availability

The code for training the networks and for the analyses performed in this work will be made available

## Data availability

All data used in the present study will be deposited as MATLAB-formatted data in Open Science Framework, https://osf.io/dqy3g/.

