## Supplementary Figures and Information for "Random noise promotes slow heterogeneous synaptic dynamics important for robust working memory computation"

### Extended data

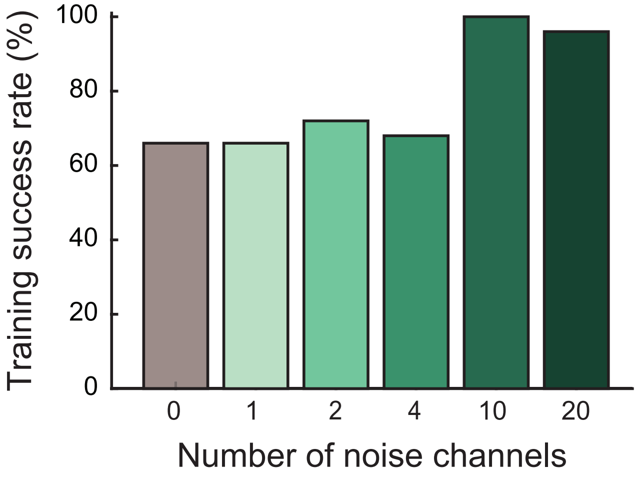


**Extended Data Fig. 1: Comparison of the training success rate on the one-delay DMS task between RNNs trained without noise and RNNs trained with varying amount of internal noise.** The grey bar is the baseline model (RNNs trained without any noise). The number of noise channels and variance of the noise signals were varied (described in Supplementary Appendix S4).

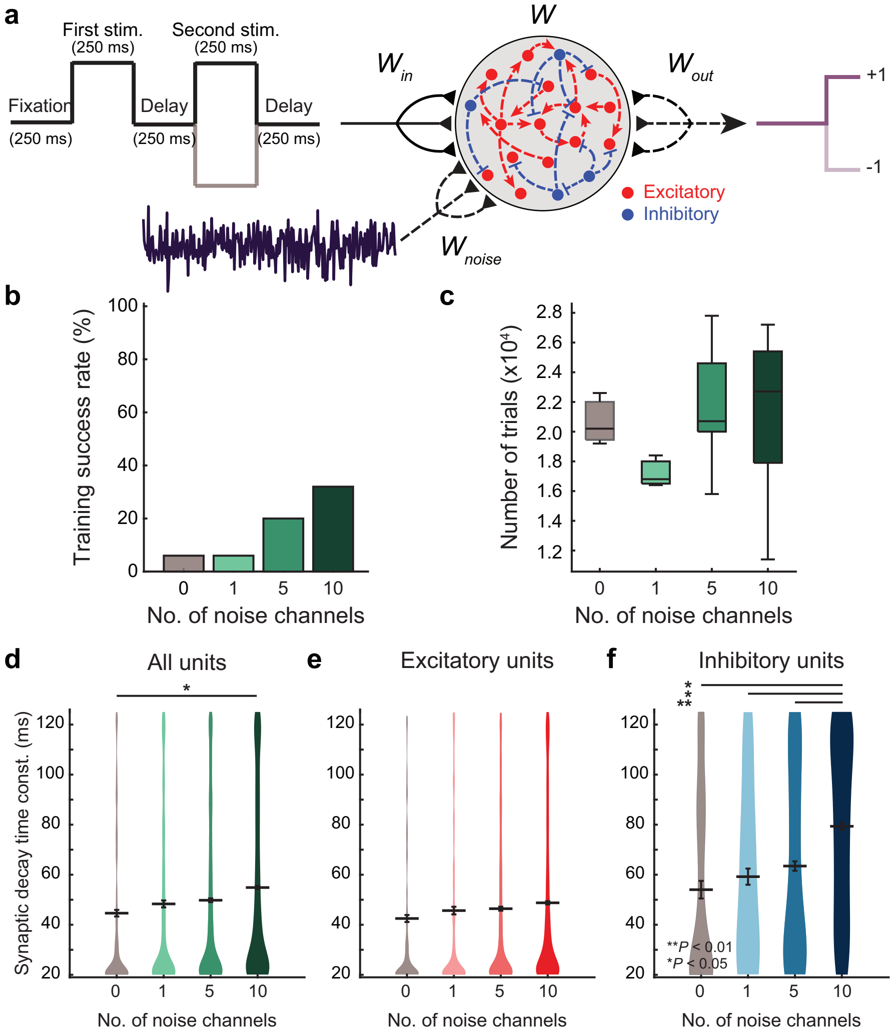


**Extended Data Fig. 2: Delayed match-to-sample (DMS) task with two delays and model dynamics.** **a**, A schematic diagram illustrates the paradigm used to trained our RNN model on a delayed match-to-sample (DMS) where two delay intervals were present. **b**, Training performance of the RNN models on the DMS task with two delay intervals. RNN models with varying amount of internal noise (i.e., 0, 1, 5, and 10 noise channels) were trained to perform this task. Training success rate was measured as the number of successfully trained RNNs (out of 50 RNNs). **c**, The average number of trials required to reach the training criteria. Boxplot: central lines, median; bottom and top edges, lower and upper quartiles; whiskers, 1.5 × interquartile range; outliers are not plotted. **d-e**, Comparison of synaptic decay time constants of RNN models trained on the DMS with two delay intervals. For each noise condition, synaptic decay time constants of successfully trained models are reported for all units (**d**), and separately for excitatory (**e**) and inhibitory units (**f**). Similar patterns regarding the effects of internal noise on the inhibitory dynamics are observed on the 2-delay DMS task as compared to the 1-delay DMS task (see Fig. 3) for all units ($P < 0.05$; $H = 9.44$; Kruskal-Wallis test with Dunn's post hoc test) and inhibitory population ($P < 0.001$; $H = 18.71$; Kruskal-Wallis test with Dunn's post hoc test). Gray horizontal lines, mean.

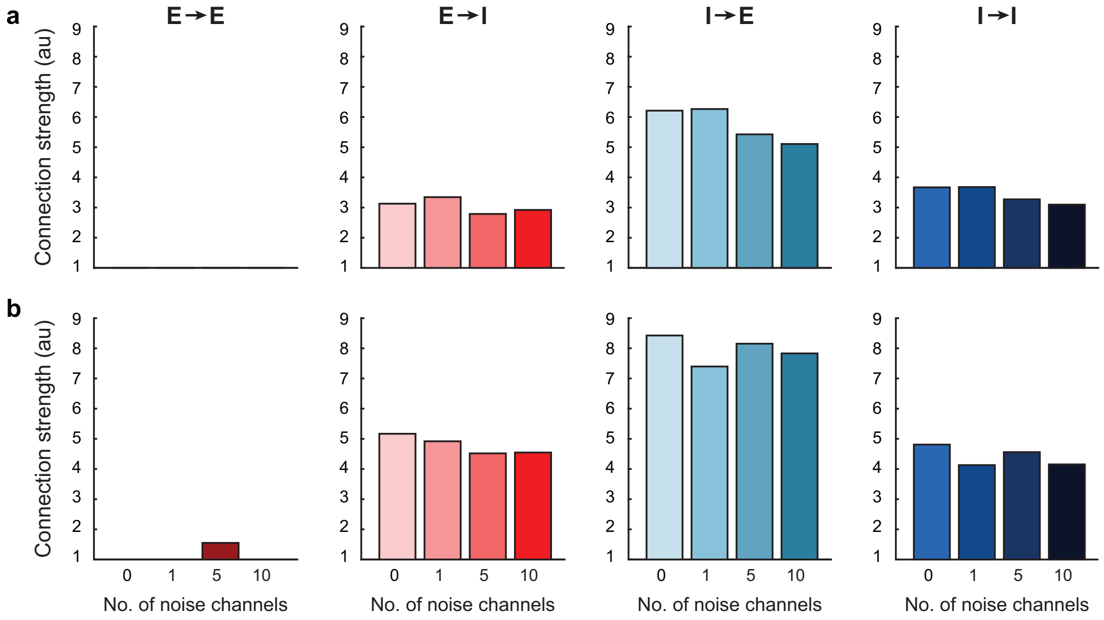


**Extended Data Fig. 3: Comparison of the average synaptic connection strength across the four noise conditions ($C = 0, 1, 5, 10$).** **a**, Average synaptic strength separated by the four synaptic connection types ($E \to E$, $E \to I$, $I \to E$, $I \to I$) for the RNNs trained to perform the DMS task. **b**, Average synaptic strength separated by the four synaptic connection types for the RNNs trained to perform the DMS task in which two delay intervals were presented.

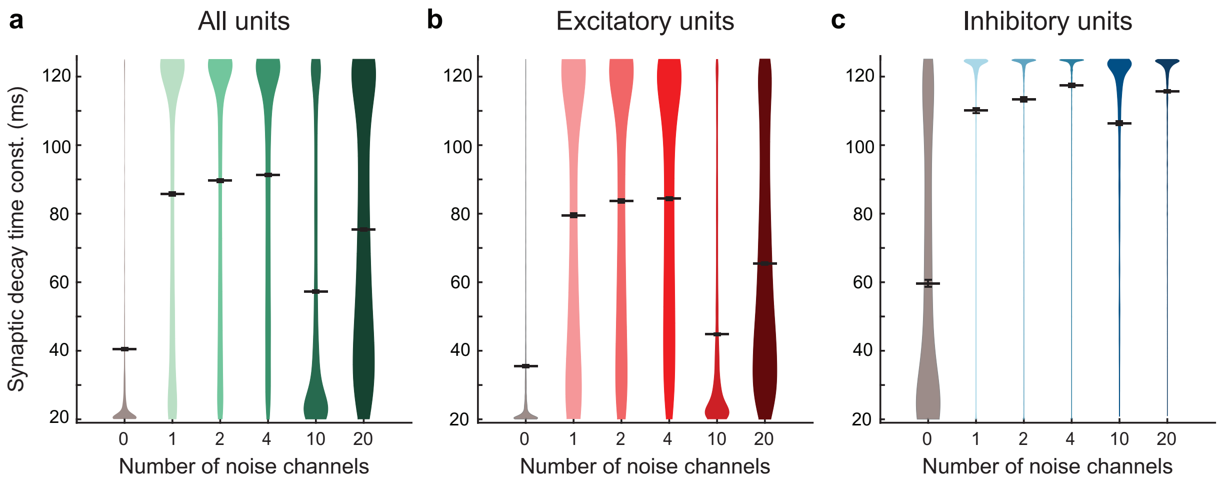


**Extended Data Fig. 4: Comparison of the synaptic decay time constant values between RNNs trained without noise and RNNs trained with varying amounts of internal noise**. The trained synaptic decay values are shown for all (**a**), excitatory (**b**), and inhibitory (**c**) units from the RNNs with different amount of internal noise structures as manipulated through the number of noise channels and variance of the noise signals (described in Supplementary Appendix S4). Gray horizontal lines, mean

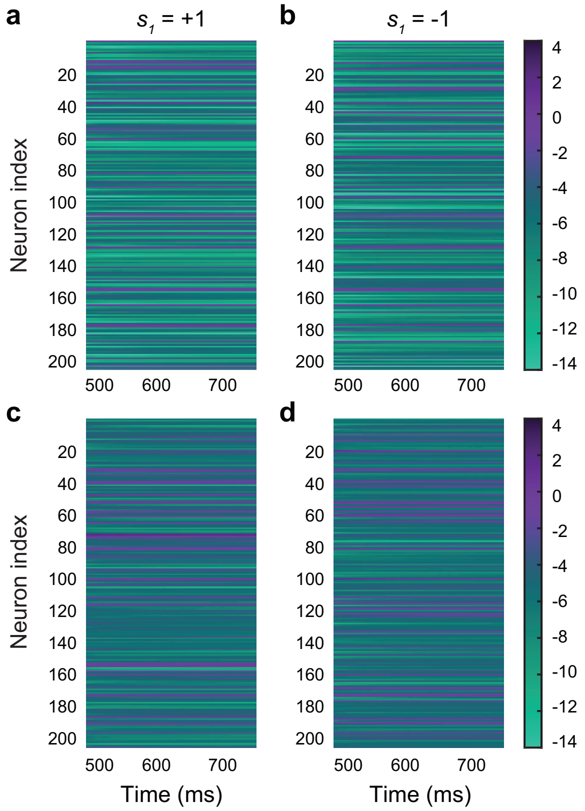


**Extended Data Fig. 5: Stable steady state activity during the delay period.** Firing rate activity ($\boldsymbol{r}$) during the delay period of the DMS task with a delay of 250 ms. The firing activity shown is of individual units within a sample RNN model that was successfully trained without noise (**a** and **b**) and with noise ($C = 10$; **c** and **d**). For both conditions, we observed stable steady states for each stimulus condition. These observed steady states during the delay interval enabled us to perform linear stability analysis during this time period to investigate the influence of noise on the maintenance of working memory representation.

### Supplementary Information

#### S1 Appendix.

**Gradient loss with respect to the synaptic decay time constant.** In this section, we derive the gradient of the loss function with respect to the synaptic decay time constant parameters ($\boldsymbol{\tau}$). First, we define the loss function ($\mathcal{L}$) as

$$\mathcal{L} = \sum_{t=1}^{T} (z(t) - o(t))^2 \tag{11}$$

$$= \sum_{t=1}^{T} (z(t) - (\boldsymbol{w}^{(\mathrm{out})}\boldsymbol{r}(t) + b))^2. \tag{12}$$

We have the following gradient of the loss function at time $t$ (i.e., $\mathcal{L}_t$) with respect to $\tau_i$ (the synaptic decay time constant for unit $i$):

$$\frac{\partial \mathcal{L}_t}{\partial \tau_i} = \frac{\partial \mathcal{L}_t}{\partial r_t^{(i)}} \frac{\partial r_t^{(i)}}{\partial x_t^{(i)}} \frac{\partial x_t^{(i)}}{\partial \tau_i}. \tag{13}$$

Note that we have denoted the component $i$ of a vector $\boldsymbol{v}$ as $v^{(i)}$ instead of $v_i$, for some vector $\boldsymbol{v}$ whose subscript labels the time step, to avoid an overcrowded notation. For $\frac{\partial \mathcal{L}_t}{\partial r_t^{(i)}}$, we first rewrite the loss function,

$$\begin{aligned} \mathcal{L}_t &= z_t - \left( \sum_{j=1}^{N} w_j^{(\mathrm{out})} r_t^{(j)} + b \right)^2 \\ &= z_t - \left( w_i^{\mathrm{out}} r_t^{(i)} + \sum_{\substack{j=1 \\ j \neq i}}^{N} w_j^{(\mathrm{out})} r_t^{(j)} + b \right)^2. \end{aligned}$$

Then we have

$$\frac{\partial \mathcal{L}_t}{\partial r_t^{(i)}} = (-2) \cdot w_i^{(\mathrm{out})} \cdot \left( w_i^{(\mathrm{out})} r_t^{(i)} + \sum_{\substack{j=1 \\ j \neq i}}^{N} w_j^{(\mathrm{out})} r_t^{(j)} + b \right). \tag{14}$$

For $\frac{\partial r_t^{(i)}}{\partial x_t^{(i)}}$, we use Equation (6) to get

$$\frac{\partial r_t^{(i)}}{\partial x_t^{(i)}} = \sigma(x_t^{(i)}) \cdot (1 - \sigma(x_t^{(i)})). \tag{15}$$

Lastly, we substitute $\alpha = \frac{\Delta}{\tau_i}$ into the discretized equation (Equation (8)) for $\frac{\partial x_t^{(i)}}{\partial \tau_i}$ to obtain

$$x_t^{(i)} = (1-\alpha)x_{t-1}^{(i)} + \alpha(\boldsymbol{w}\boldsymbol{r}_{t-1} + \boldsymbol{w}^{(\text{noise})}\boldsymbol{\psi}_{t-1} + \boldsymbol{w}^{(\text{in})}\boldsymbol{u}_{t-1}) + \xi_{t-1}^{(i)}.$$

Differentiating the above equation with respect to $\alpha$ leads to

$$\begin{aligned}\frac{\partial x_t^i}{\partial \alpha} &= \frac{\Delta}{\tau_i^2}x_{t-1}^{(i)} - \frac{\Delta}{\tau_i^2}(\boldsymbol{w}\boldsymbol{r}_{t-1} + \boldsymbol{w}^{(\text{noise})}\boldsymbol{\psi}_{t-1} + \boldsymbol{w}^{(\text{in})}\boldsymbol{u}_{t-1}) \\ &= \frac{\Delta}{\tau_i^2}(x_{t-1}^{(i)} - \boldsymbol{w}\boldsymbol{r}_{t-1} - \boldsymbol{w}^{(\text{noise})}\boldsymbol{\psi}_{t-1} - \boldsymbol{w}^{(\text{in})}\boldsymbol{u}_{t-1}).\end{aligned}$$

Since $\partial\alpha/\partial\tau_i = -\Delta/\tau_i^2$,

$$\frac{\partial x_t^{(i)}}{\partial \tau_i} = -\frac{\Delta^2}{\tau_i^4}(x_{t-1}^{(i)} - \boldsymbol{w}\boldsymbol{r}_{t-1} - \boldsymbol{w}^{(\text{noise})}\boldsymbol{\psi}_{t-1} - \boldsymbol{w}^{(\text{in})}\boldsymbol{u}_{t-1}). \tag{16}$$

Multiplying the three equations (Equations (14) to (16)) leads to the gradient of the loss function with respect to the synaptic decay parameter.

### S2 Appendix.

**Parameter initialization.** In this section, we discuss how we initialized all the parameters in our model.

We initialized each synaptic decay time constant parameter ($\tau_i$) using $\tau_i = \sigma(\mathcal{N}(0,1))\tau_{step} + \tau_{min}$, where $\sigma(\cdot)$ is the sigmoid function and $\mathcal{N}(0,1)$ refers to the standard normal distribution. $\tau_{min} =$ 20 ms and $\tau_{step} = 105$ ms were used to constrain the parameter to range from 20 ms to 125 ms.

The recurrent connectivity matrix ($\boldsymbol{w} \in \mathbb{R}^{N\times N}$) was initialized as a sparse random matrix drawn from $\mathcal{N}(0, 1.5/\sqrt{N \cdot P_c})$ with the initial connectivity probability ($P_c$) of 0.20.

The "noise" input matrix ($\boldsymbol{w}^{(\text{noise})} \in \mathbb{R}^{N\times C}$) was initialized as a full random matrix drawn from the standard normal distribution (i.e., $\mathcal{N}(0,1)$). The noise signals with $C \times T$ independent components ($\boldsymbol{\psi} \in \mathbb{R}^{C\times T}$) were initialized in the same manner. The input weight matrix ($\boldsymbol{w}^{(\text{in})} \in \mathbb{R}^{N\times U}$) was initialized as a random matrix drawn from the standard normal distribution. The external noise ($\boldsymbol{\xi}$) and the readout weights ($\boldsymbol{w}^{(\text{out})}$) were initialized as random matrices drawn from $\mathcal{N}(0, 0.01)$. The bias term ($b$ in Equation (7)) was initialized to 0.

### S3 Appendix.

**Comparison of the excitatory and inhibitory connection strength.** In order to demonstrate that adding internal noise did not lead to significant changes in the recurrent connectivity structures, we averaged the recurrent synaptic weights separated by the four connection types for each noise condition ($C = 0, 1, 5, 10$): excitatory-to-excitatory ($E \rightarrow E$), excitatory-to-inhibitory ($E \rightarrow I$), inhibitory-to-excitatory ($I \rightarrow E$), and inhibitory-to-inhibitory ($I \rightarrow I$). We analyzed the models trained to perform the one-delay DMS task (Extended Data Fig. 2a) and two-delay DMS task (Extended Data Fig. 2b). All the models, regardless of the noise condition, contained sparse $E \rightarrow E$ recurrent connections (Extended Data Fig. 2a and b). Furthermore, we did not observe any significant changes in the mean synaptic strength across the four noise conditions, suggesting that the robustness conferred by the internal noise was not due to changes in synaptic connection strength.

**S4 Appendix.**

**Comparison of different structures of internal noise.** In this section, we show the effects of different noise structures on the training success rate and synaptic decay time constants. Particularly, we used the following configurations:

- 1 channel of noise signals (i.e., $C = 1$) drawn from $\mathcal{N}(0, 400)$
- 2 channels of noise signals, each drawn from $\mathcal{N}(0, 100)$
- 4 channels of noise signals, each drawn from $\mathcal{N}(0, 25)$
- 10 channels of noise signals, each drawn from $\mathcal{N}(0, 4)$
- 20 channels of noise signals, each drawn from $\mathcal{N}(0, 1)$

For each configuration, we trained 50 RNNs to perform the one-delay DMS task described in the main text. As shown in Extended Data Fig. 3, the noise structure with high variance ($C = 1, 2, 4$) did not provide any improvement in the training success rate compared to the RNNs trained without any noise ($C = 0$). Adding more than 10 noise channels ($C = 20$) did not result in additional improvement in the success rate (darkest green in Extended Data Fig. 3).

For the high-variance conditions ($C = 1, 2, 4$), the synaptic decay constant values for both excitatory and inhibitory were prolonged (Extended Data Fig. 4). Consistent with our main findings (Fig. 3), adding any amount of noise led to a significant increase in the inhibitory synaptic decay parameter (Extended Data Fig. 4c). These results demonstrate both the number and variance of the internal noise signals can significantly impact the optimization of the synaptic decay parameter.
